# Calcium-activated chloride channels, rather than the voltage-gated ones, regulate sperm capacitation

**DOI:** 10.64898/2026.09.29.755310

**Authors:** Adeel Ahmad, Jesús Martínez-Hernández, Carine Corcini, Ayazhan Akthatova, Karolina Nowicka-Bauer, Elisabeth Pinart, Marc Yeste

**Affiliations:** Biotechnology of Animal and Human Reproduction (TechnoSperm), Institute of Food and Agricultural Technology, University of Girona, ES-17003 Girona, Spain; Unit of Cell Biology, Department of Biology, Faculty of Sciences, University of Girona, ES-17003 Girona, Spain; Department of Cell Biology and Histology, Faculty of Medicine, IMIB-Pascual Parrilla, International Excellence Campus for Higher Education and Research “Campus Mare Nostrum”, University of Murcia, ES-30120 Murcia, Spain; Comparative Animal Reproduction, Department of Animal Pathology, College of Veterinary Medicine, Federal University of Pelotas, Brazil

**Keywords:** Sperm, capacitation, acrosome reaction, voltage-gated chloride channels, calcium-activated chloride channels

## Abstract

**Background:** Ion channels regulate mammalian sperm function, yet chloride channels have received less attention than their cation counterparts. Herein, we investigated the presence, localization, and function of voltage-gated chloride channels (ClC-1, ClC-2, and ClC-K) and calcium-activated channels (ANO1 and ANO2) during sperm capacitation and the acrosome reaction.

**Results:** Using the pig as an animal model, we localized these channels in different sperm regions: ClC-1 in the equatorial and post-acrosomal regions; ClC-2 in the connecting and principal pieces; ClC-K in the acrosome, connecting, and principal pieces; ANO1 in the principal piece; and ANO2 in the mid-piece and post-acrosomal sheath region. Following this, we addressed the roles of these channels in sperm capacitation and acrosome reaction by blocking them with different inhibitors: 9AC, specific for ClCs; BBR, specific for CaCCs; and NPPB, a broad-spectrum blocker that inhibits ClCs, CaCCs, and other chloride channels. Blocking CaCCs with BBR led to a significant reduction in sperm motility and mitochondrial membrane potential, with effects similar to those of the non-selective inhibitor NPPB or to a combination of BBR and NPPB. Moreover, although none of the three inhibitors affected intracellular calcium levels or acrosomal integrity, a significant reduction in protein tyrosine phosphorylation was observed in sperm treated with the combination of NPPB and BBR. Finally, membrane lipid disorder was increased when ClCs were blocked with 9AC and when sperm were incubated with a combination of BBR and NPPB.

**Conclusions:** Our results suggest that CaCCs play a major role in sperm capacitation compared to ClCs. Further research to address the regulatory mechanisms involved and the potential relationship with other ion channels is warranted.

## 1. Introduction

In eutherian mammals, the ejaculated spermatozoon is unable to fertilize the oocyte and rather needs to undergo a series of biochemical and structural changes within the female reproductive tract known as capacitation [1]. Sperm capacitation was first described by Austin and Chang in the 1950s and involves, among others, changes in motility patterns, with capacitated sperm displaying hyperactivated movement; cholesterol efflux and an increase in plasma membrane lipid disorder; rise in intracellular calcium levels; tyrosine phosphorylation of certain sperm proteins; and pH alkalinization [2]. All these modifications are required for sperm to trigger the acrosome reaction and subsequently fertilize the oocyte [3–8].

The ability of the spermatozoon to undergo capacitation depends on several factors, including pH, plasma membrane potential, and a balanced ionic environment [9,10]. In this context, ion channels are relevant in regulating the modifications that occur during the transport throughout the male reproductive system to the female reproductive tract, including epididymal maturation, activation of motility, capacitation, and the acrosome reaction [11–14]. Ion channels allow the facilitated diffusion of millions of ions across the plasma membrane and, along with ion pumps, are crucial for the regulation of intracellular pH, hyperpolarization, cell volume, and calcium [15–18].

Protons, calcium, potassium, sodium, and bicarbonate are among the ions that play vital roles in key sperm physiological events, such as motility, capacitation, and acrosomal reaction [14,17,19,20]. As compared to all these ions, the function of chloride has been investigated less, despite separate ion channels having been identified in murine, bovine, and human sperm, including: Cystic Fibrosis Transmembrane Conductance Regulator (CFTR) [21,22]; Calcium-activated Chloride Channels (CaCCs) [23]; voltage-gated Chloride Channels (ClCs) [24]; γ-aminobutyric acid (GABA)-gated, and glycine-gated receptors [25]]. ClCs were first recognized in the marine ray *Torpedo marmorata* [26], and since then,, nine members of this family have been identified, each comprising 10 to 12 transmembrane segments with N- and C-termini in the cytoplasm [27,28]. CaCCs belong to the anoctamin family, which is composed of 10 members (ANO1 - ANO10) that are encoded by *TMEM16a*-*TMEM16K* genes, respectively [29–31]. Whether ClC and CaCC regulate sperm capacitation and the acrosome reaction, however, remains to be investigated.

As this study used the pig as a model, we first confirmed the presence and localization of ClCs (ClC-1, ClC-2, and ClC-K) and CaCCs (ANO1 and ANO2) in the sperm of this species. Following this, we addressed the roles of ClCs and CaCCs in sperm capacitation and progesterone-induced acrosome exocytosis by examining the effects of different inhibitors. 9-anthracenecarboxylic acid (9AC) was used to inhibit ClCs [32], benzbromarone (BBR) [33] was utilized to block CaCCs (Hwang et al., 2016)(Centeio et al., 2020), and 5-nitro-2-(3-phenylpropylamino) benzoic acid (NPPB), which is less selective, was employed to inhibit chloride channels in general, including ClCs and CaCCs [34–36]. Based on the results of the individual inhibitors, we then conducted an additional series of experiments, in which we combined BBR and NPPB. Our hypothesis was that both ClCs and CaCCs regulate sperm capacitation and the progesterone-induced acrosomal exocytosis.

## 2. Materials and methods

### 2.1. Semen samples

A total of 96 seminal doses were used in this study, each obtained from a different boar and sourced from a local company (Semen Cardona, S.L., Barri des Segalers, Cardona, Spain) operating under standard commercial conditions. According to the company, all boars were sexually mature (18-24 months of age), healthy, housed under standardized humidity and temperature conditions, fed with a standard diet, and had free access to water. Following semen collection, ejaculates were diluted in a commercial extender (Vitasem, Magapor, S.L., Ejea de los Caballeros, Spain) to a final concentration of 33 million sperm/mL in 90-mL doses. The diluted semen was then cooled to 17 °C and transported to the University of Girona within 12 hours of collection. Animals were handled exclusively by farm staff in accordance with the guidelines established by the Regional Government of Catalonia (Barcelona, Spain). The authors were not involved in animal handling, and the seminal doses used were commercial products intended for artificial insemination. Hence, specific approval from an ethical committee was not required for this study.

### 2.2. Experimental design

Five experiments were conducted. The first experiment investigated the presence and localization of ClCs and CaCCs. The second, third and fourth experiments interrogated how inhibiting ClCs with 9AC, chloride channels as a whole with NPPB, and CaCCs with BBR, respectively, affected sperm capacitation and progesterone-induced acrosome exocytosis. Based on the results of the second, third, and fourth experiments, we conducted an additional fifth trial in which BBR and NPPB were combined.

For each biological replicate, three seminal doses obtained, each from a different boar, were pooled and centrifuged at 600 × g and 17 °C for 5 min. After centrifugation, sperm pellets were resuspended in a capacitation medium (TCM: 20 mM HEPES, 100 mM NaCl, 3.1 mM KCl, 5 mM glucose, 21.7 mM sodium L-lactate, 1 mM sodium pyruvate, 0.3 mM Na₂HPO₄, 0.4 mM MgSO₄·7H₂O, 4.5 mM CaCl₂·2H₂O, 5 mg/mL bovine serum albumin (BSA), and 15 mM sodium bicarbonate) to a final concentration of 1 × 10⁷ sperm/mL. The suspension was then divided into four aliquots. One aliquot was used as a control (without any inhibitor), whereas the other three were added with different inhibitor concentrations (9AC, BBR, NPPB, or NPPB+BBR, depending on the experiment). Additionally, another aliquot, which served as a negative control, was prepared in a non-capacitating medium (TBM: 20 mM HEPES, 100 mM NaCl, 3.1 mM KCl, 5 mM glucose, 21.7 mM sodium L-lactate, 1 mM sodium pyruvate, 0.3 mM Na₂HPO₄, and 0.4 mM MgSO₄·7H₂O) to a final concentration of 1 × 10⁷ sperm/mL without any treatment. The concentrations tested were selected following preliminary experiments, in which the effects of each inhibitor were evaluated using motility and viability. The concentrations chosen were 0.1 mM, 1 mM, and 10 mM for 9AC (experiment 2); 50 µM, 100 µM, and 200 µM for NPPB (experiment 3), 1 µM, 10 µM, and 100 µM for BBR (experiment 4); and 100 µM NPPB + 10 µM BBR, 100 µM NPPB + 100 µM BBR, and 200 µM NPPB + 100 µM BBR for experiment 5. Samples were incubated at 38 °C under 100% humidity and 5% CO₂ for 3 h (Binder GmbH, Tuttlingen, Germany). After 120 min of incubation, progesterone (10 µM) was added to each sample. Sperm capacitation-related parameters were evaluated at 0, 60, 120, 130, and 180 min of incubation using CASA and flow cytometry. Each experiment (i.e., experiment 2: 9AC; experiment 3: NPPB; experiment 4: BBR; and experiment 5: NPPB + BBR) was replicated eight times.

### 2.3. Immunoblotting

The presence of ClC and CaCC was first confirmed with immunoblotting. Briefly, 1 mL of the pooled semen samples was centrifuged at 800 × g and 4 °C for 10 min. Supernatants were discarded, and the sperm pellet was washed twice with cold phosphate-buffered saline (PBS) at 4 °C and centrifuged under the same conditions. Proteins in the pellet were extracted through resuspension in a lysis buffer (PBS supplemented with 1% SDS, 1% protease inhibitor cocktail (Sigma, Sant Luis, MO, USA), 0.1 mM phenyl-methane-sulfonylfluoride (PMSF), and 700 mM sodium orthovanadate). After thorough mixing, samples were incubated on ice for 30 min with occasional agitation, and then centrifuged at 14,000 × g and 4 °C for 20 min. Total protein in the supernatants was quantified using a detergent-compatible method with a commercial kit (Bio-Rad Laboratories, United States).

After determining protein concentration based on a bicinchoninic acid assay (BCA; ThermoFisher Scientific), samples were incubated with mixed loading buffer (Laemmli Buffer; Bio-Rad Laboratories, United States) supplemented with 10% (v/v) β-mercaptoethanol at 95 °C for 5 min. Fifteen μg of total protein was loaded per lane into a Mini-PROTEAN TGX 4-15% Bis-Tris SDS-PAGE precast gel (Bio-Rad Laboratories, United States). In the first lane of the gel, 10 μL of a Precision Plus Protein All Blue Prestained Standard (Bio-Rad Laboratories, United States) was loaded, and gel electrophoresis was run at 180 V for 45 min. Following this, proteins were transferred onto a PVDF membrane by a Trans-Blot Turbo device (1 A and 25 V for 30 min; Bio-Rad Laboratories, United States). Once proteins were transferred, membranes were blocked with TBS-Tween-20 supplemented with 5% BSA (Bovine serum albumin) at room temperature with agitation for 1 h. Membranes were incubated with the following primary antibodies (Alomone Labs, Jerusalem, Israel; dilution factors indicated between brackets, v/v): ClC-1 (1/5000; ref.: ACL-005), ClC-2 (1/1000; ref.: ACL-002), ClC-K (1/5000; ref.: ACL-004), ANO1 (1/1000; ref: ACL-011), and ANO2 (1/1000; ref: ACL-012) in TBS-T supplemented with 5% BSA at 4 °C overnight. Membranes were subsequently washed three times in TBS-T, and then incubated with a secondary HRP-coupled goat anti-rabbit antibody (Dako, Denmark) at double the dilution of the primary antibody (v/v; i.e., 1/2000 for CLC-2, ANO1, and ANO2; 1/10000 for CLC-1 and CLC-K) in TBS-T containing 5% BSA, at room temperature for 60 min. Finally, membranes were washed five times, exposed to a chemiluminescent substrate (Millipore, United States), and visualized with a G:BOX Chemi XL system (SynGene, United States). Blocking peptide assays (peptides were present at a fivefold excess relative to their primary antibodies) and negative controls (no primary antibody) were included to confirm the specificity of all primary antibodies.

### 2.4. Immunofluorescence

Immunostaining with the same antibodies used for immunoblotting was performed to localize ClC and CaCC in sperm. These experiments were conducted in duplicate, with each biological replicate consisting of a pool of three seminal doses. Samples were centrifuged at 800 × g for 5 min to remove the preservation medium. The resulting pellet was resuspended in the same volume of PBS. Samples were again centrifuged at 800 × g for 5 min, and pellets were fixed in a 4% paraformaldehyde solution (ThermoFisher, Germany) at room temperature for 20 min. Fixed samples were again centrifuged at the conditions mentioned before, and the resulting sperm pellets were resuspended in PBS at a final concentration of 5×10^6^ sperm/mL. Following this, 50 µL of each sample was smeared onto slides. Smears were either non-permeabilized (for observation of ANO1 and ANO2) or permeabilized (for observation of ClC-1, ClC-2, and ClC-K) with PBS containing 1% Triton X-100 at room temperature for 30 min, gently washed in PBS (5 min), and blocked with 0.02 M glycine in PBS at room temperature for 20 min in a humidity chamber. After that, samples were washed in PBS (5 min) and incubated with rabbit antibodies against ClC-1 (1/200; ref.: ACL-005), ClC-2 (1/200; ref.: ACL-002), ClC-K (1/200; ref. ACL-004), ANO1 (1/200; ACL-011), and ANO2 (1/100; ACL-012) in PBS containing 1% BSA (v/w), at 4 °C overnight in a humidity chamber. The next day, slides were washed in PBS three times (5 min each) and incubated with a secondary antibody in PBS containing 1% BSA (v/w) at 4 °C overnight in a humidity chamber protected from light. For ClC-1, ClC-2, ClC-K, and ANO1, this antibody was a secondary donkey anti-rabbit antibody conjugated to Alexa Fluor 488 (Invitrogen, United States) at a dilution of 1/400 (v/v). In the case of ANO2, this antibody was an Alexa Fluor 647-conjugated secondary antibody, at a dilution of 1/200 (v/v). Finally, smears were washed five times in PBS and mounted with one drop of antifading mounting medium (ProLong™ Gold Antifade Mountant; ThermoFisher Scientific) containing DAPI. Samples were kept in the dark before examination under a confocal laser-scanning microscope (CLSM, Nikon A1R; Nikon, Tokyo, Japan).

The specificity of primary antibodies was, like for immunoblotting, confirmed by pre-adsorption with its blocking peptide (fivefold excess) at room temperature for 1 h. Negative controls were performed by omitting the primary antibody.

### 2.5. Determination of sperm motility

Sperm motility was evaluated using a computer-assisted sperm analysis (CASA) system consisting of a phase-contrast microscope (Olympus BX41; Olympus, Tokyo, Japan), a video camera, a computer, and a sperm-specific image analysis software (ISAS; V1.0; Proiser, S.L.; Valencia, Spain). Five µL from each sample was placed onto a prewarmed Makler Chamber (Sefi Medical Instruments; Haifa, Israel) and observed under a phase-contrast microscope with a 10× objective (negative phase-contrast). A total of 1000 sperm per sample were captured and analyzed by ISAS software. Percentages of total motile and progressively motile sperm (%) were recorded for each sample, along with the following kinetic parameters: curvilinear velocity (VCL, µm/s), straight-line velocity (VSL, µm/s), average-path velocity (VAP, µm/s), linearity (LIN, %), straightness (STR, %), wobble (WOB, %), amplitude of lateral head displacement (ALH, µm), and beat-cross frequency (BCF, Hz). Sperm with a VAP equal to or greater than 10 μm/s were considered motile, and those with a STR equal to or greater than 45% were considered progressively motile.

### 2.6. Flow cytometry

Mitochondrial membrane potential, reactive oxygen species, membrane lipid disorder, acrosome integrity, intracellular calcium levels, and tyrosine phosphorylation were evaluated using a CytoFlex Flow Cytometer (Beckman Coulter; Brea, CA, USA). Each parameter was determined following staining with a specific combination of fluorochromes purchased from Thermo Fisher Scientific (Waltham, MA, USA). Details of such evaluations are presented according to MIFlowCyt guidelines [37].

All samples were excited with a blue laser (488 nm), except those stained with Live/Dead Fixable Far Red (LD), which were excited with a 638-nm laser. The FITC filter (524/40) was used for Yo-Pro-1, PNA-FITC, Fluo4, JC-1 monomers (JC-1_mon_), and 2’,7’-dichlorofluorescein (DCF) fluorochromes. The PE filter (585/42) was utilized to detect ethidium (E), JC-1 aggregates (JC-1_agg_), and merocyanine 540 (M540) fluorochromes. APC (660/20) and PC5.5 (690/50) filters were employed for Alexa Fluor647-conjugated anti-pTyr antibody and propidium iodide (PI), respectively. The sperm population was identified using Forward Scatter (FSD) and Side Scatter Detectors (SSD), which analyze particle size and roughness, respectively.

#### 2.6.1. Mitochondrial membrane potential

Mitochondrial membrane potential was determined with the metachromatic fluorochrome JC-1 (5,5’,6,6’-Tetrachloro-1,1’,3,3’-tetraethylbenzimidazolylcarbocyanine iodide). At low mitochondrial membrane potential, JC-1 remains in its monomeric state (JC-1_mon_) and emits green fluorescence with a peak at 520 nm. When mitochondrial membrane potential is high, JC-1 molecules form aggregates (JC-1_agg_) and emit red fluorescence with a peak at 596 nm. Sperm were incubated with JC-1 (final concentration: 750 nM) and LIVE/DEAD (LD) fixable far red fluorochromes (diluted 1:8000 in PBS) at 38 °C for 30 min [38]. The ratio of JC-1_agg_ to JC-1_mon_ in viable sperm (LD^-^) was measured as mitochondrial membrane potential.

#### 2.6.2. Intracellular levels of reactive oxygen species (ROS)

Total levels of reactive oxygen species (ROS) were evaluated with 2’,7’-dichlorodihydrofluorescein diacetate (H_2_DCFDA) and PI [39]. H₂DCFDA is a non-fluorescent molecule that is oxidized by reactive oxygen species (ROS) to dichlorofluorescein (DCF), which emits green fluorescence. Briefly, samples were incubated with H₂DCFDA (50 µM) at 38 °C for 20 min, and then with PI (6 µM) for 5 min under the same conditions. The geometric mean fluorescence intensity (GeoMean) of DCF in viable sperm (PI^-^) and total sperm was measured as an indicator of ROS levels.

#### 2.6.3. Intracellular levels of superoxides

Superoxide (O_2_^-^) levels in sperm were determined by co-staining the samples with 5 µM of HE and 31.25 nM of Yo-Pro-1, following the protocol of Guthrie and Welch [39]. HE permeates the plasma membrane of sperm and is oxidized by superoxide ions to ethidium (E), which results in the emission of red fluorescence. Samples were incubated with HE (5 µM) and Yo-Pro-1 (31.25 nM) at 38 °C for 20 min in the dark. Superoxide levels were evaluated as geometric mean fluorescence intensity (GeoMean) of E in viable sperm (YP^-^) and total sperm.

#### 2.6.4. Membrane lipid disorder

Lipid disorder of the sperm plasma membrane was determined following the protocol of Rathi et al. [40], which is based on Merocyanine 540 (M540). When the lipid disorder of the plasma membrane increases, M540 intercalates into that membrane and emits red fluorescence, whereas Yo-Pro-1 (YP) labels sperm with an increased membrane permeability. Based on these markers, four sperm populations were identified: (1) viable sperm with low membrane lipid disorder (M540^−^/YP^−^), (2) viable sperm with high membrane lipid disorder (M540^+^/ YP^−^), (3) non-viable sperm with low membrane lipid disorder (M540^−^/YP^+^), and (4) non-viable sperm with high membrane lipid disorder (M540^+^/YP^+^). Briefly, samples were incubated with M540 (2.5 µM) and Yo-Pro-1 (25 nM) at 38 °C for 10 min in the dark. Results are expressed as geometric mean fluorescence intensity (GeoMean) of M540 in viable sperm (YP^-^) and the total sperm population. In addition, the percentages of viable (YP^-^) and non-viable sperm (YP^+^) with low (M540^-^) and high (M540^+^) membrane disorder were also calculated.

#### 2.6.5. Acrosome integrity

Acrosomal integrity was determined using the Cooper and Yeung protocol with few modifications [41]. Samples were stained with the Live/Dead fluorochrome at 38 °C in the dark for 20 min. Samples were then centrifuged at 1000 × g and room temperature for 3 min, resuspended in a fixation solution (10 mL PBS containing 4% PFA, 10 µL Triton, and 3 µL T-20), and incubated at room temperature for 60 min. After incubation, samples were centrifuged again at 1000 × g for 3 min, and the pellets were resuspended in 200 µL of PBS. PNA-FITC (1.17 µM) was added to the resuspended samples, which were then incubated at 38 °C in the dark for 15 min. Results were expressed as the geometric mean fluorescence intensity (GeoMean) of PNA in viable and total sperm.

#### 2.6.6. Intracellular Calcium levels

To evaluate intracellular Ca^2+^ levels, sperm were double-stained with Fluo4-AM and PI [42]. Fluo4-AM, upon entering the cell, de-esterifies to Fluo4 and, if present, binds to calcium, emitting green fluorescence. Thus, the intensity of green fluorescence emitted by Fluo4 is directly proportional to intracellular calcium levels. Samples were incubated with Fluo4 (1.17 µM) and PI (5.6 µM) at 38 °C in the dark for 10 min . Results are expressed as the geometric mean fluorescence intensity (GeoMean) of Flou4 in viable sperm (PI^-^).

#### 2.6.7. Tyrosine-phosphorylation of sperm proteins

Tyrosine phosphorylation (pTyr) analysis was conducted by the protocol established by Peris-Frau et al. with slight modifications [43]. First, samples were incubated with Live/Dead solution at 38 °C in the dark for 20 min. Subsequently, samples were centrifuged at 1,000× g and room temperature for 3 min, resuspended in 10 mL blocking buffer (5% BSA in PBS), incubated for 1 min, and centrifuged again under the same conditions. Sperm pellets were then resuspended in 4% paraformaldehyde and incubated at room temperature for 15 min. Samples were then centrifuged at 1,000× g and room temperature for 3 min, resuspended in PBS, and stored at 4 °C overnight. Samples were centrifuged at 1,000× g and room temperature for 3 min, resuspended in permeabilization buffer (0.5 g BSA, 100 μL Triton X-100, and 0.02 g sodium azide in 10 mL PBS), and incubated at room temperature for 60 min. After centrifugation at 1,000× g and room temperature for 3 min, sperm pellets were either resuspended with the antibody solution (blocking solution with anti-pTyr antibody conjugated with AlexaFluor488 (Abcam) at 1:1,000), or resuspended in blocking buffer (negative control, without the antibody). Samples were incubated together at 4 °C under agitation in the dark overnight. Samples were centrifuged at 1,000× g and room temperature for 3 min, resuspended in PBS, and analyzed using a flow cytometer. Results were expressed as fold change (relative to the negative control) of the geometric mean fluorescence intensity (GeoMean) of pTyr in both viable (LD^-^) and total sperm populations.

### 2.7. Statistical analyses

Statistical analyses were conducted using R (Ver. 4.52; R Foundation for Statistical Computing, Vienna, Austria) within the RStudio integrated development environment (ver. 2026.01.1+403 “Apple Blossom”). Data were preprocessed using readxl, dplyr, tidyr, purrr, stringr, forcats, and tibble packages. Testing of assumptions was carried out with the car package, and statistical models were built with afex, lme4, lmerTest, and glmmTMB. The emmeans package was used for calculating the estimated marginal means and making pairwise comparisons. Broom and broom.mixed packages were used for model outputs.

In all experiments, the treatment (which varied in each case: 9AC, BBR, NPBB) was the between-subjects factor and the incubation time was the within-subjects factor, with subject identity being treated as a random effect to account for within-subject correlations. Analyses followed a hierarchical model-selection strategy based on model assumptions. Normality of residuals and homogeneity of variance were tested through the Shapiro-Wilk and Levene tests, respectively. When data matched with parametric assumptions (i.e., normality and homoscedasticity), they were analyzed using repeated-measures analysis of variance (rmANOVA) with type III sums of squares using the afex package with sum contrasts. When deviations from normality were detected, appropriate data transformations were applied, and the data were analyzed using linear mixed-effects models (LMMs). If Gaussian assumptions could not be met or the response variable exhibited strictly positive skewed distributions, generalized linear mixed models (GLMMs) with a Gamma distribution were fitted using glmmTMB. In cases where none of the above approaches were suitable, linear mixed models were used on raw data as a fallback. Post-hoc pairwise comparisons were conducted using estimated marginal means with Tukey adjustment for multiple comparisons.

Results are presented as estimated means ± standard error (SEM), and statistical significance was set at *P* ≤ 0.05. Graphics were generated using the ggplot2 package, and significant differences between treatments within a given time point were displayed with different letters, as provided by the multcompView package.

## 3. Results

### 3.1. ClC-1, ClC-2, and ClC-K are present in pig sperm and have different localization patterns

Immunoblotting confirmed the presence of ClC-1, ClC-2, and ClC-K in pig sperm. Specifically, a band with an apparent molecular weight of approximately 120 kDa was observed for CLC-1. According to the UniProt database (entry: F1SRV8; species: *Sus scrofa*), ClC-1 in pigs has 986 amino acids and its molecular weight is 108,788 Da (Fig. 1a). For ClC-2, two bands with apparent molecular weights of approximately 65 kDa and 95 kDa were detected (Fig. 1b). As indicated in the UniProt database (entry: A0A480UNX6; species: *Sus scrofa*), ClC-2 in pigs has 905 amino acids and a molecular weight of 99,203 Da. Finally, a single band of approximately 90 kDa was observed for ClC-K (Fig. 1c). According to the UniProt database (entry: F1SUU0; species: Sus scrofa), ClC-K in pigs has 644 amino acids and a molecular weight of 70,367 Da. The specificity of the primary antibody was confirmed in all cases by negative controls and blocking peptide assays (Suppl. Fig. 1a-c).

**Fig. 1.**
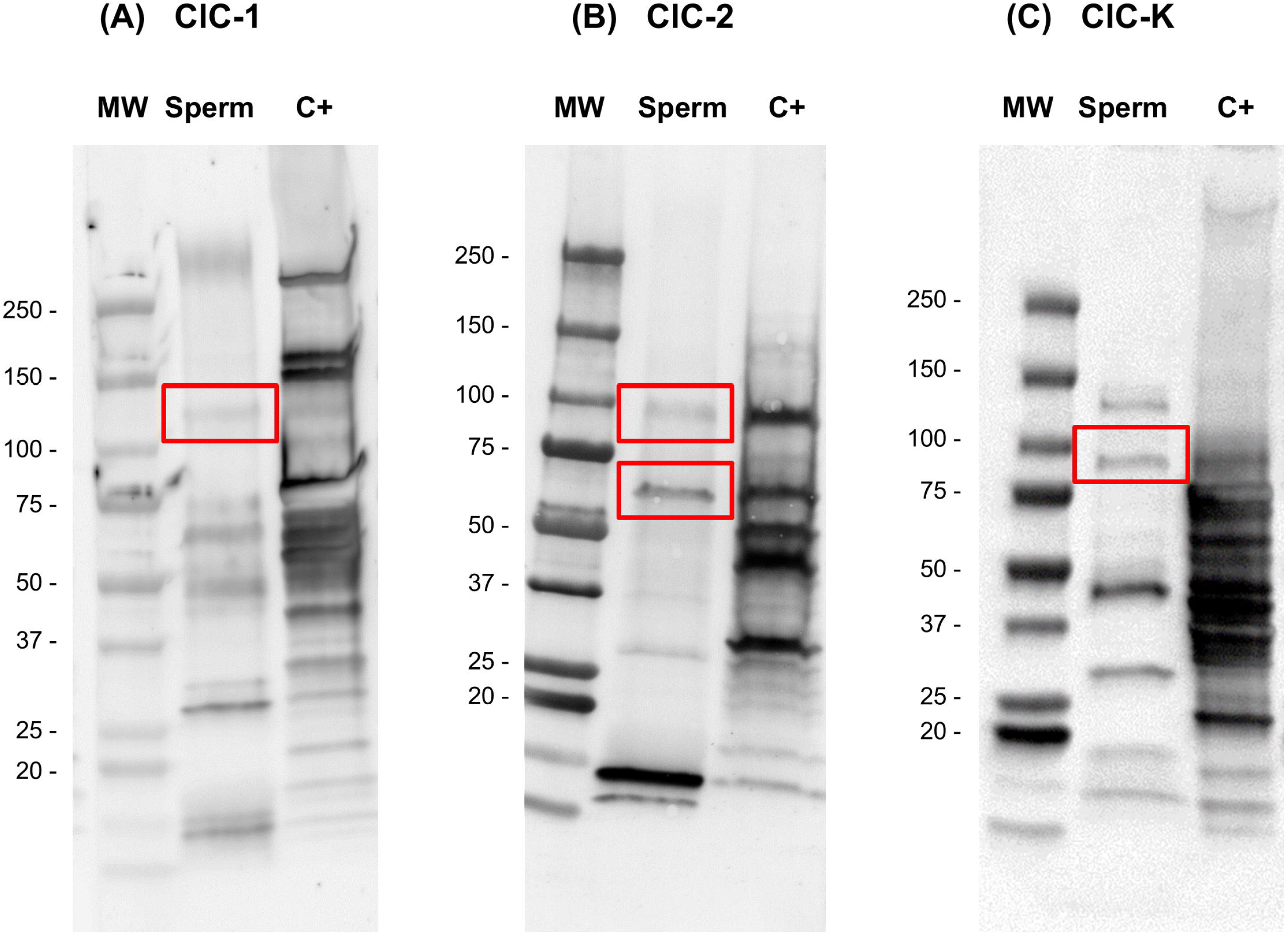
Immunoblotting of voltage-gated chloride channels (ClC-1, ClC-2, and ClC-K) in pig sperm. (A) Incubation with the anti-ClC-1 antibody revealed a single band at ∼120 kDa. C+: pig kidney; (B) incubation with the anti-ClC-2 antibody revealed two bands at ∼ 65 kDa and ∼ 95 kDa. C+: pig brain; and (C) incubation with the anti-ClC-K antibody revealed one band at ∼ 85 kDa. C+: pig kidney. MW: molecular weight ladder.

Immunofluorescence revealed that ClC-1 is localized in the post-acrosomal region and the equatorial segment of sperm (Fig. 2c). In contrast, staining of ClC-2 (Fig. 2f) showed strong labeling in the principal piece and the connecting piece (neck), as well as weak labeling in the midpiece of the sperm. Finally, ClC-K exhibited a heterogeneous labeling pattern, typically observed in the acrosome, connecting piece (neck), and principal piece (Fig. 2i). Yet, some sperm lacked ClC-K, as evidenced by the absence of staining for this protein (Figure 2c). As in the previous case, the specificity of primary antibodies was confirmed by blocking peptide assays (Suppl. Fig. 2) and negative controls (i.e., incubation with secondary but without primary antibody).

**Fig. 2.**
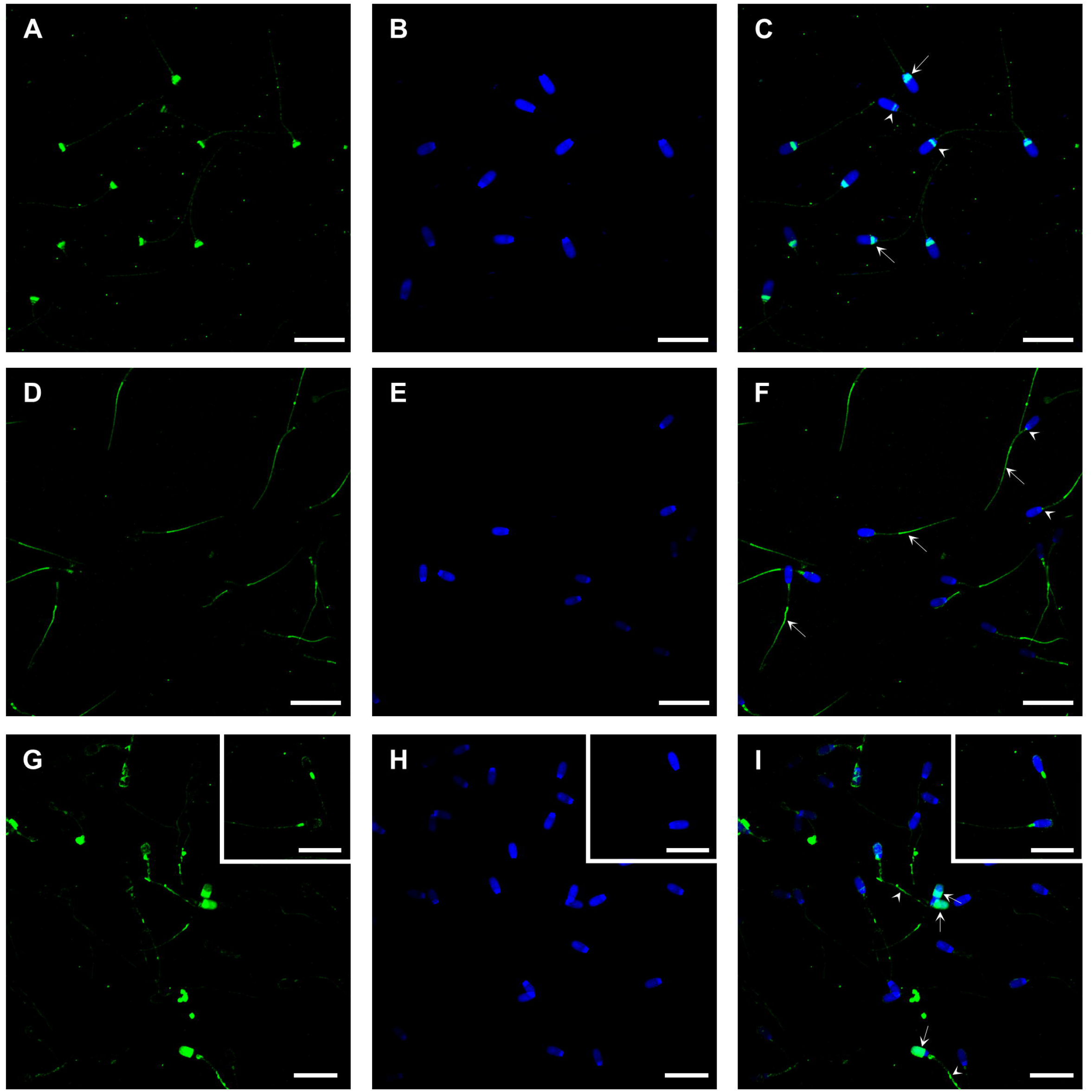
Immunolocalization of voltage-gated chloride channels (ClC-1, ClC-2, and ClC-K) in pig sperm through confocal laser scanning microscopy. (A-C) CLC-1. (A) Immunostaining for ClC-1 in the green channel. (B) DAPI staining in the blue channel. (C) Merge of the previous channels showing positivity in the post-acrosomal region (arrow) and the pre-equatorial segment (arrowhead). (D–F) Immunostaining for ClC-2. (D) Immunostaining for ClC-2 in the green channel. (E) DAPI staining in the blue channel. (F) Merge of the previous channels showing positivity in the principal piece (arrow) and the connecting piece or neck (arrowhead). (G–I) Immunostaining for ClC-K. (G) Immunolabeling for ClC-K in the green channel. (H) DAPI staining in the blue channel. (I) Merge of the previous channels showing positivity in the acrosome (arrow), principal piece (arrowhead), and connecting piece or neck (inset detail). Scale bar: 20 µm.

### 3.2. CaCCs are also present in pig sperm

Immunoblotting revealed specific bands for both ANO1 and ANO2. In the case of ANO1, a specific band heavier than 100 kDa but lighter than 150 kDa was observed (Fig. 3a). As, according to the Uniprot database (entry: I3LJG3; species: *Sus scrofa*), this protein has 1,005 amino acids and its molecular weight is 114,886 Da, it is likely that this band corresponds to the protein. The blocking peptide assay demonstrated the specificity of the primary antibody (Suppl. Fig. 3a). Regarding ANO2, we observed a clear band at approximately 75 kDa and another at 150 kDa. According to the Uniprot database (entry: A0A5G2QQ54; species: *Sus scrofa*), this protein has 997 amino acids and its molecular weight is 113,340 Da (Fig. 3b). As blocking peptide experiments (Suppl. Fig. 3b) demonstrated the specificity of the primary antibody, it could be that the lighter band corresponds to a degraded form of the protein, and the heavier band corresponds to a glycosylated form of the protein, as this protein is known to have different glycosylation sites.

**Fig. 3.**
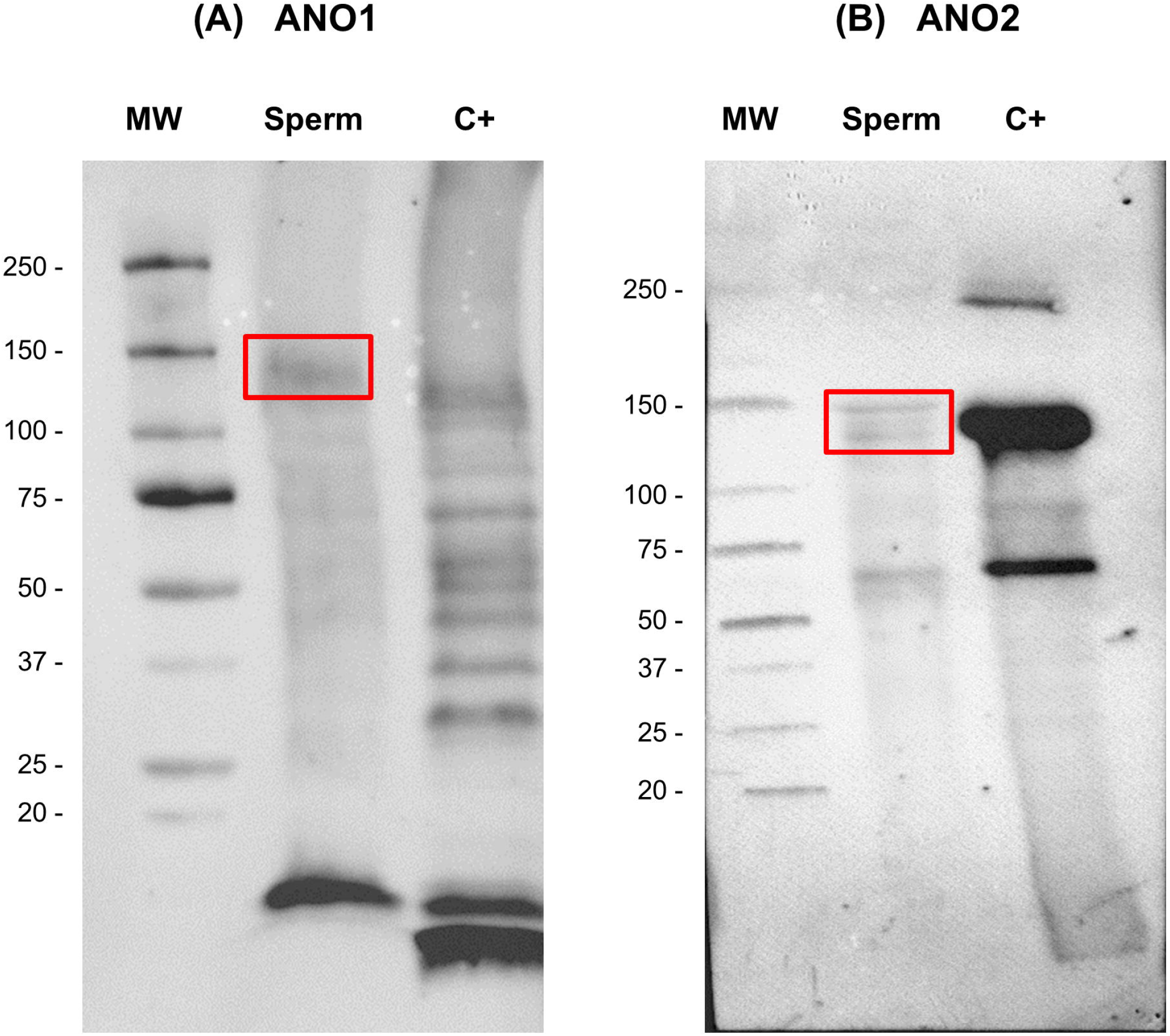
Immunoblotting of calcium-activated chloride channels (ANO1 and ANO2) in pig sperm. (A) Incubation with the anti-ANO1 antibody revealed a single band at ∼130-140 kDa. C+: pig kidney; (B) incubation with the anti-ANO2 antibody revealed two bands at ∼ 65 kDa and ∼ 130-150 kDa. C+: pig kidney. MW: molecular weight ladder.

Immunofluorescence analyses showed that ANO1 in sperm is mainly distributed along the sperm tail, with stronger fluorescence observed in the principal piece and discrete staining in the head region (Fig. 4c). There were some sperm cells, however, that did not display labeling for ANO1. Moreover, ANO2 was mainly found in the tail, with stronger staining in the mid-piece and some staining in the post-acrosomal sheath region of the head (Fig. 4f). The specificity of the primary antibodies was demonstrated by both blocking peptide assays (Suppl. Fig. 4) and negative controls (i.e., incubation with secondary but without primary antibody).

**Fig. 4.**
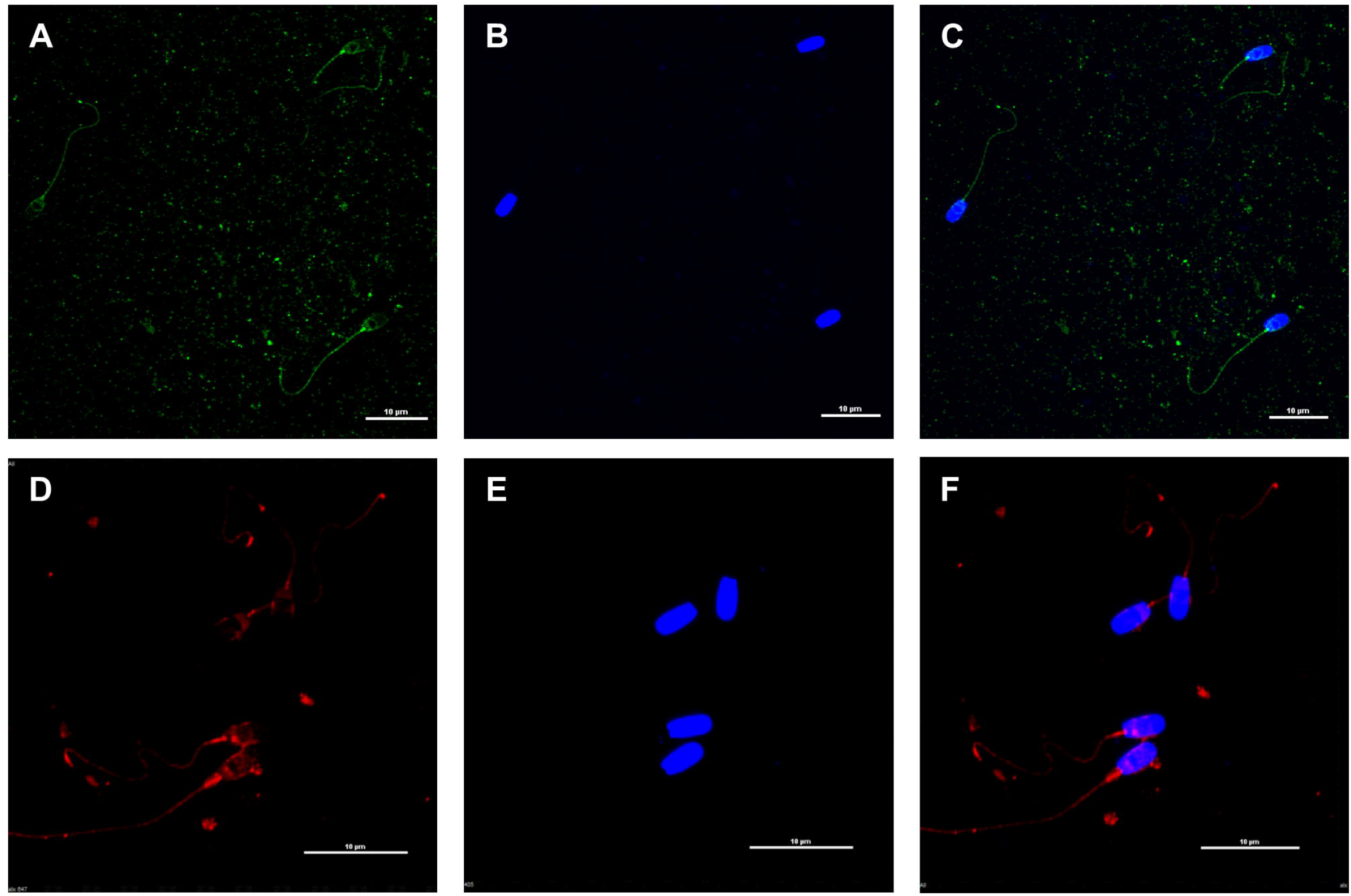
Immunolocalization of calcium-activated chloride channels (ANO1 and ANO2) in pig sperm through confocal laser scanning microscopy. (A-C) ANO1. (A) Immunostaining for ANO1 in the green channel. (B) DAPI staining in the blue channel. (C) Merge of the previous channels showing stronger fluorescence in the principal piece and discrete staining in the head region. (D–F) Immunostaining for ANO2. (D) Immunostaining for ANO2 in the red channel. (E) DAPI staining in the blue channel. (F) Merge of the previous channels showing stronger staining in the mid-piece and some staining in the post-acrosomal sheath region. Scale bar: 10 µm.

### 3.3. CaCCs are involved in the regulation of sperm motility during capacitation

Inhibiting CaCCs with 100 µM BBR decreased total motility compared to the control after 130 min and 180 min of incubation (Fig. 5c). These results were similar to those observed when the general inhibitor NPPB (200 µM) was used to block chloride channels as a whole, with a decrease in total motility after 130 min (Fig. 5b), and in progressive motility after 130 min and 180 min of incubation (Fig. 6b). The combination of NPPB with BBR gave similar results, as 100 µM NPPB + 100 µM BBR significantly decreased progressive motility after 130 min of incubation (Fig. 6d), and total motility tended to be lower than the control after 60, 130, and 180 min of incubation (Fig. 5d). On the other hand, blocking ClCs with 1 mM or 10 mM 9AC also resulted in lower total (Fig. 5a) and progressive sperm motility (Fig. 6a), but these differences were not statistically significant.

**Fig. 5.**
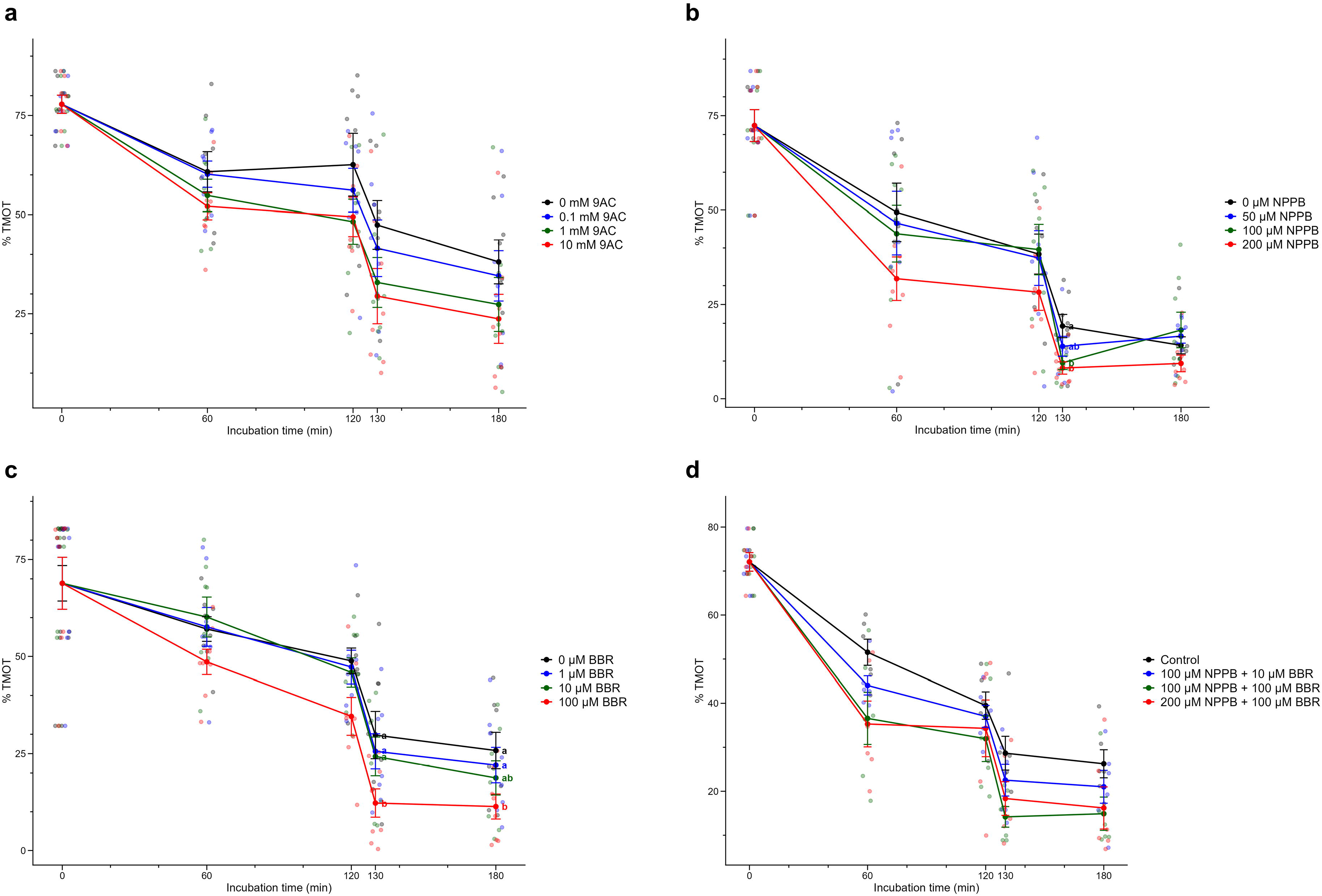
Effects of inhibiting **(a)** ClCs with 9AC (0, 0.1, 1, and 10 mM); **(b)** chloride channels as a whole with NPPB (0, 50, 100, and 200 µM), or **(c)** CaCCs with BBR (0, 1, 10, and 100 µM), and **(d)** of treating sperm with NPPB + BBR (control, 100 µM NPPB + 10 µM BBR, 100 µM NPPB + 100 µM BBR, and 200 µM NPPB + 100 µM BBR) on total sperm motility, during capacitation and progesterone-induced acrosome exocytosis. Samples were examined at 0, 60, 120, and 180 min of incubation at 38 °C during the capacitation experiment. Different letters (a, b) indicate significant differences between treatments at a given time point. Results are expressed as mean ± SEM (n = 8).

**Fig. 6.**
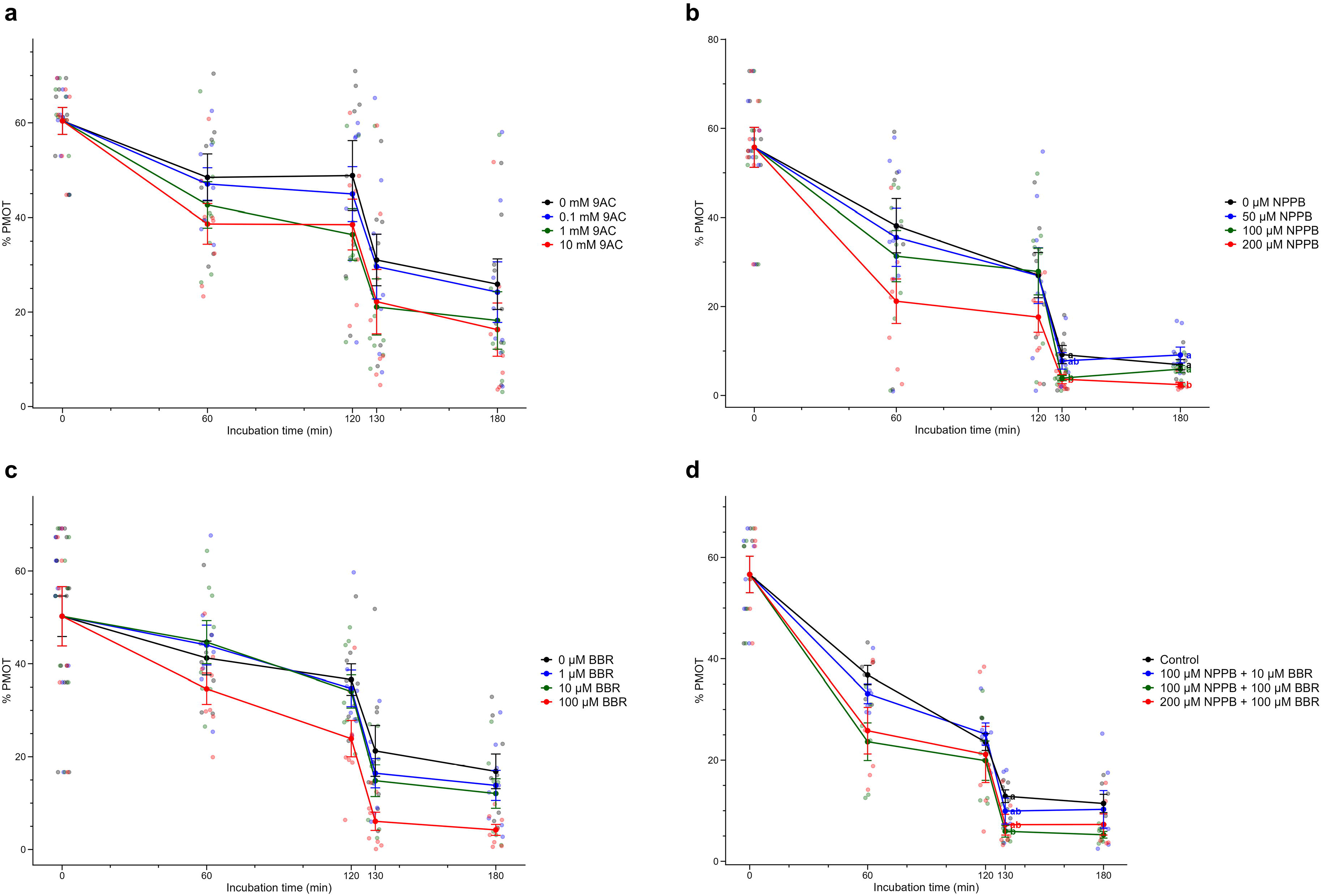
Effects of inhibiting **(a)** ClCs with 9AC (0, 0.1, 1, and 10 mM); **(b)** chloride channels as a whole with NPPB (0, 50, 100, and 200 µM), or **(c)** CaCCs with BBR (0, 1, 10, and 100 µM), and **(d)** of treating sperm with NPPB + BBR (control, 100 µM NPPB + 10 µM BBR, 100 µM NPPB + 100 µM BBR, and 200 µM NPPB + 100 µM BBR) on progressive sperm motility, during capacitation and progesterone-induced acrosome exocytosis. Samples were examined at 0, 60, 120, and 180 min of incubation at 38 °C during the capacitation experiment. Different letters (a, b) indicate significant differences between treatments at a given time point. Results are expressed as mean ± SEM (n = 8).

Inhibiting chloride channels with 200 µM NPPB significantly reduced VSL after 60, 120, 130, and 180 min of incubation, and VAP at 60 min and 120 min (Table 1). Similarly, the inhibition of CaCCs with 100 µM BBR reduced VSL after 120 and 130 min of incubation, and VAP after 130 and 180 min of incubation (Table 1). In contrast, neither NPPB or BBR had any effect on VCL, nor did blocking ClCs with 9AC on VCL, VSL, or VAP (Table 1).

Blocking chloride channels with 200 µM NPPB, but not CaCCs with BBR, decreased STR at 120 min of incubation and ALH after 180 min of incubation (Table 2). Consistent with the effects of NPPB alone, the combination of 200 µM NPPB + 100 µM BBR significantly decreased STR after 130 min and 180 min of incubation. The effects of blocking CaCCs with 100 µM BBR and blocking chloride channels with 200 µM NPPB had similar effects on BCF, as did treating sperm with 100 µM NPPB + 100 µM BBR and with 200 µM NPPB + 100 µM BBR, which decreased BCF after 130 min and 180 min of incubation (Table 2). In contrast, neither LIN nor WOB was affected by the inhibition of chloride channels and CaCCs with NPPB or BBR, respectively (Suppl. Table 1).

Finally, no kinematics parameter (VCL, VSL, VAP, LIN, STR, WOB, BCF, or ALH) was altered when ClCs were inhibited with 9AC (Tables 1 and 2; Suppl. Table 1). This would indicate that the CaCCs are the main chloride channels regulating sperm motility during capacitation, yet the involvement of other chloride channels distinct from ClCs should not be ruled out, given the effects of NPPB.

### 3.4. Inhibiting CaCCs but not ClCs decreases mitochondrial activity

Inhibition of CaCCs but not ClCs altered MMP, as shown in Fig. 7. While blocking ClCs with 9AC did not affect the JC1_agg_/JC1_mon_ ratio in viable sperm (Fig. 7a), this variable decreased, in a dose-dependent manner, when CaCCs were blocked with BBR, with lower JC1_agg_/JC1_mon_ ratios after 60 min and 180 min of incubation (Fig. 7c). At 60 min, the highest concentration of BBR (100 µM) significantly decreased MMP compared with the other treatments (0, 1, and 10 µM BBR). Similarly, at 180 min, 10 µM and 100 µM concentrations showed a significant reduction in MMP compared to the control (0 µM BBR) (Fig. 7c). The general inhibition of chloride channels with NPPB yielded similar results, as, at 60 min, all NPPB treatments decreased MMP compared with the control, with the highest concentrations (100 µM and 200 µM) showing a significantly greater reduction (Fig. 7b). The effect of BBR and NPPB on CaCCs was consistent with the effects observed when sperm were treated with 100 µM NPPB + 100 µM BBR and 200 µM NPPB + 100 µM BBR, which led to lower MMP compared to the control after 60 min of incubation (Fig. 7d). This suggests that the CaCCs are the main chloride channels regulating mitochondrial activity during sperm capacitation.

**Fig. 7.**
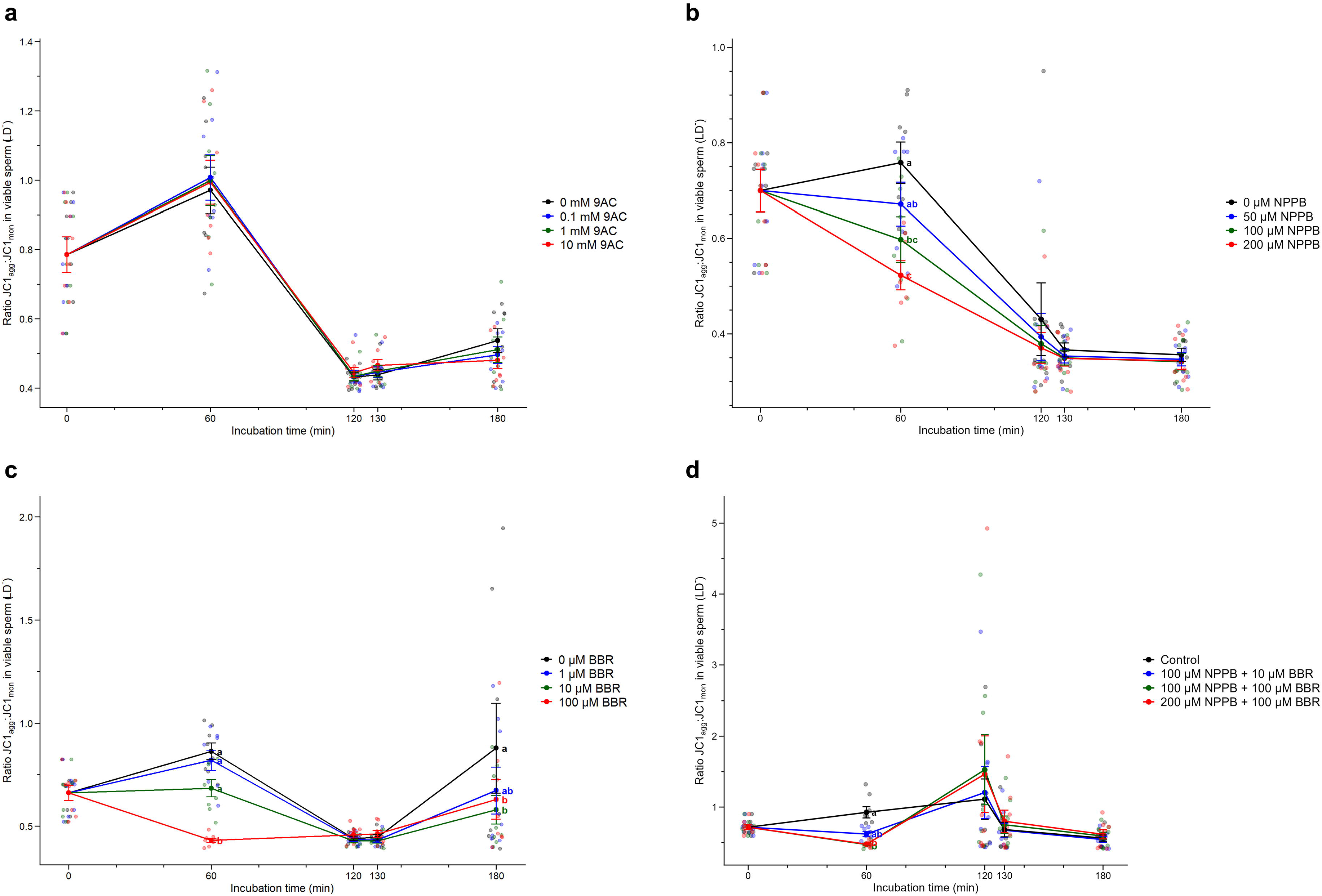
Effects of inhibiting **(a)** ClCs with 9AC (0, 0.1, 1, and 10 mM); **(b)** chloride channels as a whole with NPPB (0, 50, 100, and 200 µM), or **(c)** CaCCs with BBR (0, 1, 10, and 100 µM), and **(d)** of treating sperm with NPPB + BBR (control, 100 µM NPPB + 10 µM BBR, 100 µM NPPB + 100 µM BBR, and 200 µM NPPB + 100 µM BBR) on mitochondrial membrane potential in viable sperm, measured as ratio JC1_agg_:JC1_mon_ in the viable sperm population (LD^-^). Samples were examined at 0, 60, 120, and 180 min of incubation at 38 °C during the capacitation experiment. Different letters (a, b) indicate significant differences between treatments at a given time point. Results are expressed as mean ± SEM (n = 8).

### 3.5. Inhibition of CaCCs with BBR increases superoxide levels after 180 min of incubation

Blocking CaCCs with 100 µM BBR was observed to increase superoxide levels in viable sperm, after 180 min of incubation (Fig. 8c). In contrast, inhibiting ClCs with 9AC had no effect on superoxide levels (Fig. 8a), nor did blocking chloride channels with the general inhibitor NPPB (Fig. 8b) or incubating sperm with a combination of BBR and NPPB (Fig. 8d), even in treatments containing 100 µM BBR. Yet, a trend for 100 µM NPPB + 100 µM BBR and 200 µM NPPB + 100 µM BBR to present higher superoxide levels in viable sperm than the control was observed after 120 min and 180 min of incubation, but these differences were not statistically significant (Fig. 8d).

**Fig. 8.**
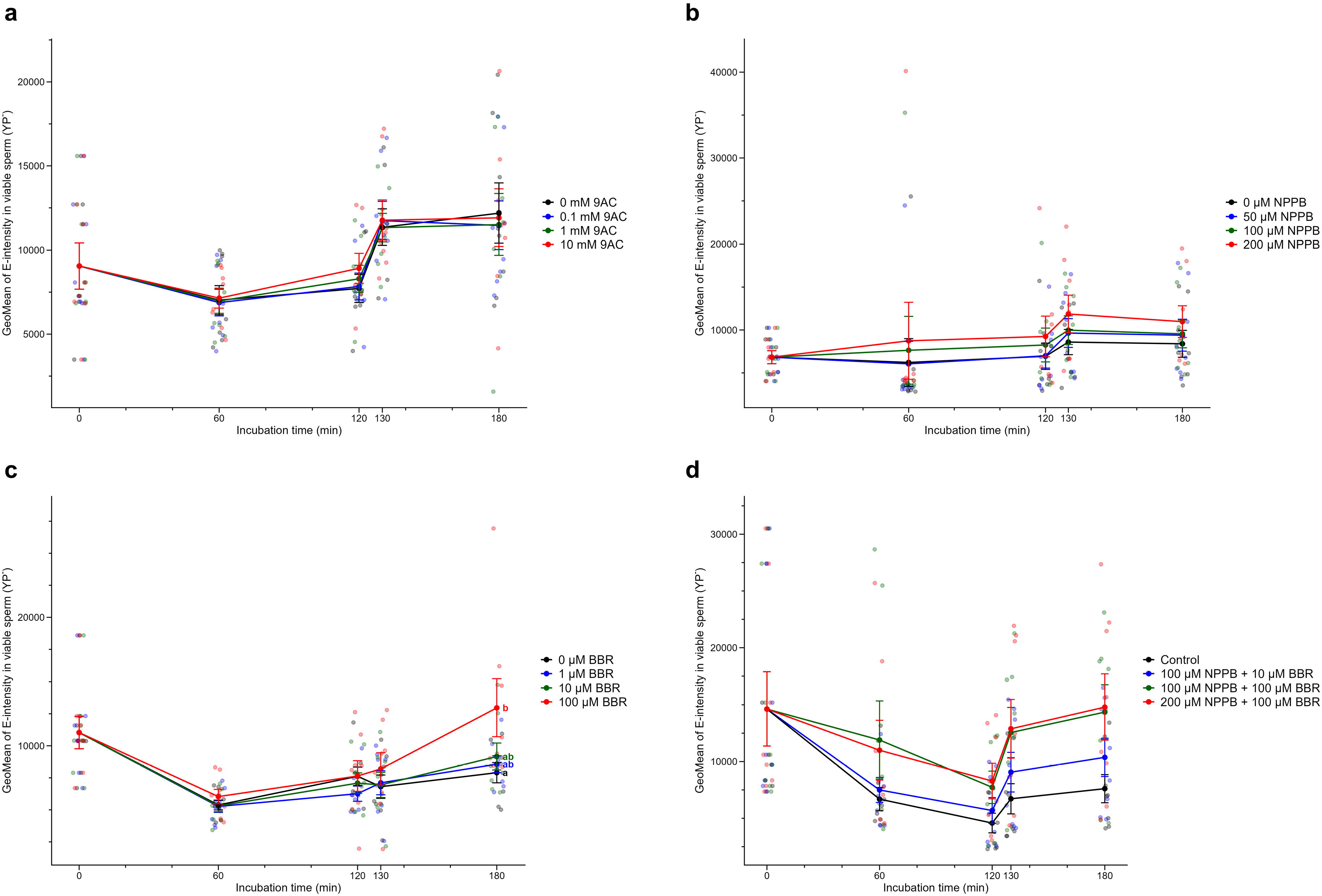
Effects of inhibiting **(a)** ClCs with 9AC (0, 0.1, 1, and 10 mM); **(b)** chloride channels as a whole with NPPB (0, 50, 100, and 200 µM), or **(c)** CaCCs with BBR (0, 1, 10, and 100 µM), and **(d)** of treating sperm with NPPB + BBR (control, 100 µM NPPB + 10 µM BBR, 100 µM NPPB + 100 µM BBR, and 200 µM NPPB + 100 µM BBR) on superoxide levels, measured as the geometric mean (GeoMean) of E intensity, in the viable sperm population (YP^-^). Samples were examined at 0, 60, 120, and 180 min of incubation at 38 °C during the capacitation experiment. Different letters (a, b) indicate significant differences between treatments at a given time point. Results are expressed as mean ± SEM (n = 8).

In contrast, blocking chloride channels in general, or specifically ClCs or CaCCs, with the aforementioned inhibitors did not result in changes in the total ROS levels in viable sperm (Suppl. Fig. 5). Noticeably, it seemed that sperm treated with 10 mM 9AC, which is a blocker of ClCs, increased total ROS levels in viable sperm, but this effect was not statistically significant (Suppl. Fig. 5a).

### 3.6. Inhibiting ClCs with 9AC increases membrane lipid disorder during sperm capacitation

Inhibiting ClCs with 1 mM and 10 mM 9AC increased membrane lipid disorder in total sperm, evaluated as the GeoMean of M540 in total sperm, after 60, 120, and 180 min of incubation (Fig. 9a). Blocking CaCCs with BBR or chloride channels in general did not alter membrane lipid disorder in total sperm (Fig. 9b and 9c), but this parameter increased at 60 min when these channels were inhibited with 100 µM NPPB + 100 µM BBR or 200 µM NPPB + 100 µM BBR (Fig. 9d), suggesting a potential effect of chloride channels other than CaCCs. In contrast, when considering only the viable sperm population, no inhibitor altered membrane lipid disorder (Suppl. Fig. 6).

**Fig. 9.**
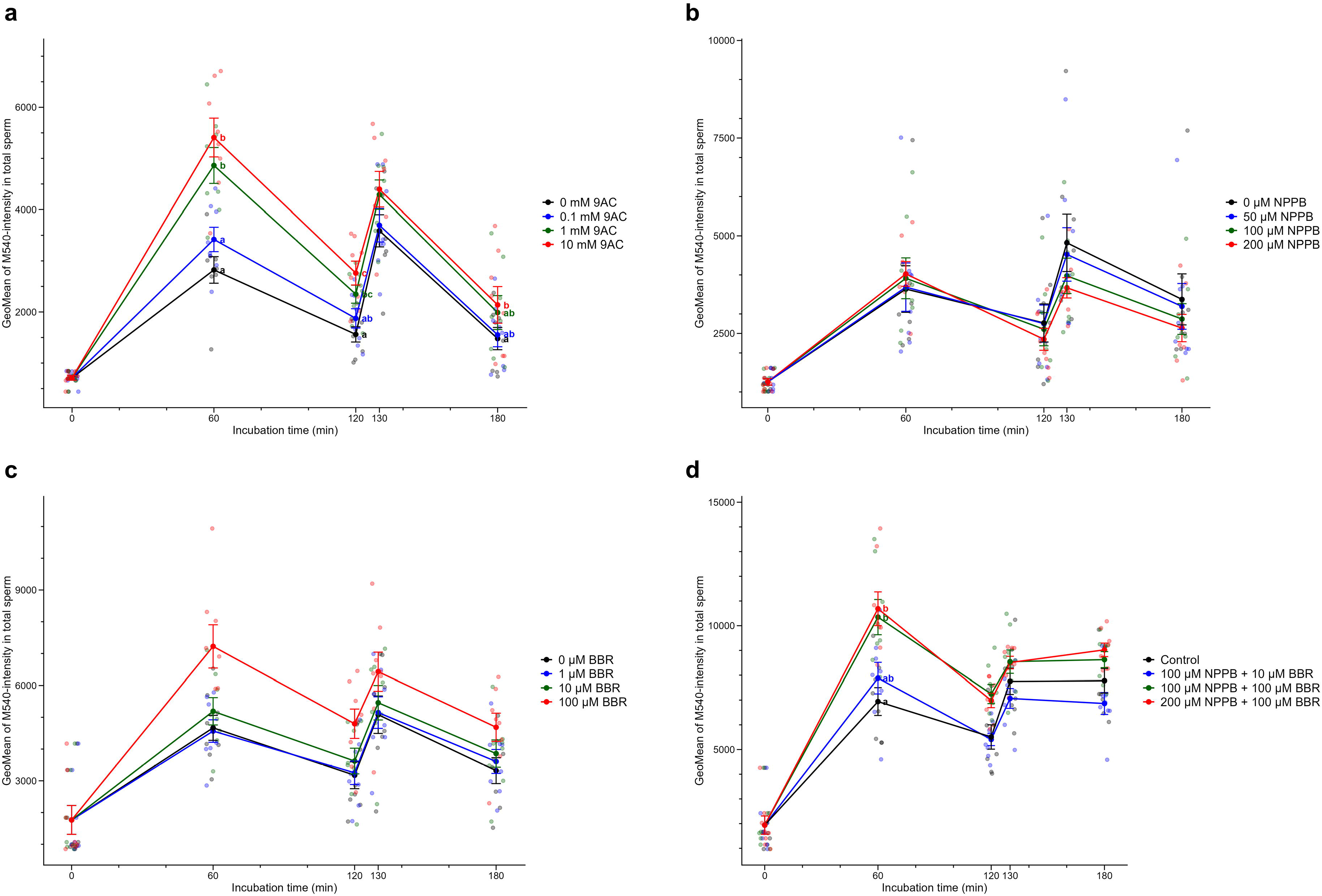
Effects of inhibiting **(a)** ClCs with 9AC (0, 0.1, 1, and 10 mM); **(b)** chloride channels as a whole with NPPB (0, 50, 100, and 200 µM), or **(c)** CaCCs with BBR (0, 1, 10, and 100 µM), and **(d)** of treating sperm with NPPB + BBR (control, 100 µM NPPB + 10 µM BBR, 100 µM NPPB + 100 µM BBR, and 200 µM NPPB + 100 µM BBR) on membrane lipid disorder, measured as the geometric mean (GeoMean) of M540 intensity in total sperm, during capacitation and progesterone-induced acrosome exocytosis. Samples were examined at 0, 60, 120, and 180 min of incubation at 38 °C during the capacitation experiment. Different letters (a, b) indicate significant differences between treatments at a given time point. Results are expressed as mean ± SEM (n = 8).

In agreement with these results, the percentages of viable sperm with low membrane lipid disorder (M540^-^/YP^-^) were significantly lower when ClCs were inhibited with 1 and 10 mM 9AC, after 60, 120, 130, and 180 min of incubation (Suppl. Fig. 7a), but did not differ from the control when chloride channels as a whole or CaCCs were blocked with NPPB or BBR, respectively, or with a combination of both (Suppl. Fig. 7b, 7c, and 7d). Similarly, the percentages of non-viable sperm with high membrane lipid disorder (M540^+^/YP^+^) were higher in samples treated with 1 mM and 10 mM 9AC than in the control after 60 and 120 min of incubation (Suppl. Fig. 8a). Furthermore, while inhibition of CaCCs with BBR or that of chloride channels with NPPB did not alter this parameter significantly (Suppl. Fig. 8b and 8c), treating samples with 100 µM NPPB + 100 µM BBR resulted in an increase in the percentage of non-viable sperm with high membrane lipid disorder at 60 min compared to the control (Suppl. Fig. 8d).

### 3.7. Blocking CaCCs reduces the levels of Tyr-phosphorylation in sperm proteins, but the effects depend on the inhibitor

Inhibiting CaCCs with 1 µM and 10 µM BBR decreased the level of pTyr in sperm proteins after 60 min of incubation (Fig. 10c). Also, treating samples with NPPB + BBR resulted in reduced levels of Tyr-phosphorylation in sperm proteins after 180 min of incubation (Fig. 10d). In contrast, neither inhibiting ClCs channels with 9AC nor inhibiting chloride channels as a whole with NPPB had any effect on pTyr (Fig. 10a and 10b), suggesting that the main impact was driven by the BBR-inhibition of CaCCs. On the other hand, the aforementioned effects were not observed when the total sperm population was considered (Suppl. Fig. 9).

**Fig. 10.**
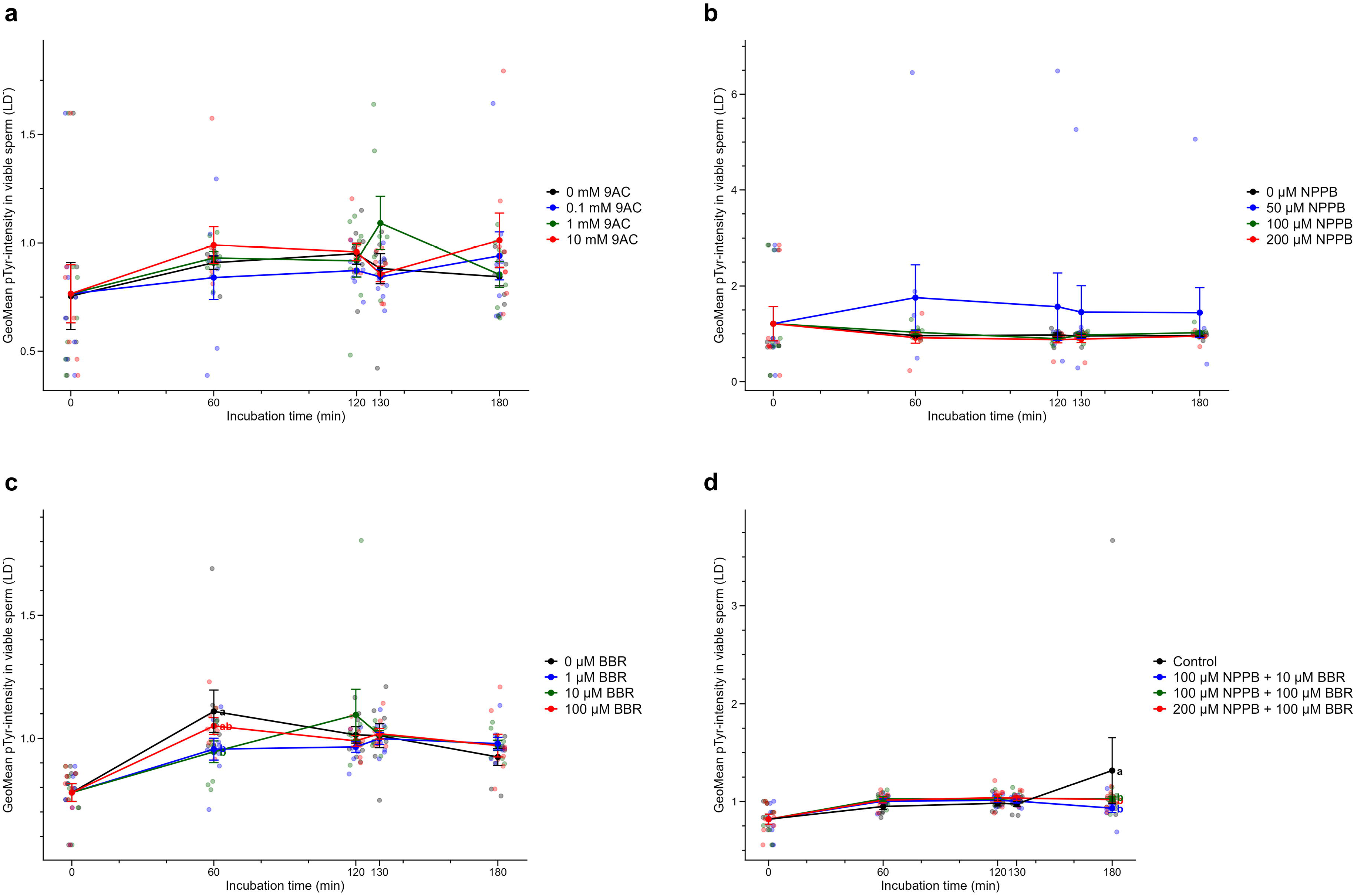
Effects of inhibiting **(a)** ClCs with 9AC (0, 0.1, 1, and 10 mM); **(b)** chloride channels as a whole with NPPB (0, 50, 100, and 200 µM), or **(c)** CaCCs with BBR (0, 1, 10, and 100 µM), and **(d)** of treating sperm with NPPB + BBR (control, 100 µM NPPB + 10 µM BBR, 100 µM NPPB + 100 µM BBR, and 200 µM NPPB + 100 µM BBR) on tyrosine phosphorylation, measured as the fold change (relative to the negative control) of geometric mean (GeoMean) of pTyr intensity in viable sperm (LD^-^). Samples were examined at 0, 60, 120, and 180 min of incubation at 38 °C during the capacitation experiment. Different letters (a, b) indicate significant differences between treatments at a given time point. Results are expressed as mean ± SEM (n = 8).

### 3.8. Blocking chloride channels has no effect on acrosome integrity or calcium levels

Inhibiting ClCs with 9AC, CaCCs with BBR, or chloride channels in general with NPPB did not have any effect on acrosome integrity, as evaluated as the GeoMean of PNA-intensity in the viable sperm population (Suppl. Fig. 10) or in the total sperm population (Suppl. Fig. 11).

Calcium levels, evaluated as GeoMean of Fluo4-intensity in total and viable sperm (Suppl. Fig. 12), were not affected by inhibiting ClC with 9AC, CaCCs with BBR, or chloride channels in general with NPPB.

## 4. Discussion

Chloride ions are considered vital for various mammalian sperm functions. During capacitation, their influx is associated with bicarbonate (HCO_3_^-^) transport, hyperactivated motility, plasma membrane hyperpolarization, and tyrosine phosphorylation [44–46]. Various chloride transporters have been investigated in mammalian sperm, such as CFTR, sodium-potassium-chloride cotransporter (NKCC), SLC26A3, and SLC26A [44,45,47]. Despite this, there is still limited information regarding the presence and localization of voltage-gated chloride channels (ClCs) and calcium-activated chloride channels (CaCCs) in mammalian sperm.

To the best of the authors’ knowledge, this is the first study assessing the presence and localization of ClCs and CaCCs in pig sperm. Through immunoblotting, we confirmed the presence of ClCs (ClC-1, ClC-2, and ClC-K) and CaCCs (ANO1 and ANO2). Regarding ClCs, we detected a band of ∼120 kDa for ClC-1, which would match the molecular weight indicated in the UniProt database for this protein (108,788 Da); two bands (∼65 kDa and ∼95 kDa) for ClC-2 (UniProt: 99,203 Da), and a band of ∼90 kDa for ClC-K (UniProt: 70,367 Da). While in some cases the molecular weights did not exactly match those reported in the UniProt database, our findings supported a higher molecular weight for ClC-1 than for ClC-2 and ClC-K. Furthermore, while two bands were observed for ClC-2, blocking peptide assays confirmed the specificity of the antibody. The reasons the molecular weights did not exactly match could be diverse, including the possibility that sperm cells express specific isoforms. As far as we are aware, no previous studies have investigated the presence of ClCs in mammalian sperm by immunoblotting, except for ClC-3 (CLCN3), which was identified in human sperm with a molecular weight of 80 kDa (Yeung et al., 2005). In human skeletal muscle cells, ClC-1 has a molecular weight of 100-150 kDa (Thomassen et al., 2018), which would also align with our finding. Moreover, ClC-2 was observed in the mouse brain and intestine with a band of approximately 90-97 kDa (Gyömörey et al., 2000), and another study involving mice testes and brain identified a cluster of specific bands ranging between 100 and 130 kDa (Göppner et al., 2021), which would agree with our results, as we saw two bands rather than one. Finally, two bands of ∼50 and 75 kDa were identified for ClC-K in protein extracts of a human airway epithelial cell line (Mummery et al., 2005). Although in our case we observed only one band for ClC-K, this suggests that, despite the predicted molecular weights in the protein database, it is not uncommon to observe more than one specific band.

In the case of CaCCs, immunoblotting revealed two bands for ANO1 (100 kDa and 150 kDa); according to UniProt, the molecular weight of this protein in pigs is 114,886 Da. These two bands were specific, as indicated by blocking peptide assays. The presence of CaCCs in sperm was previously suggested by patch-clamp recordings, as inhibitors of TMEM16A/ANO1 (NFA and DIDS) led Orta et al. to conclude that sperm expressed these channels (Orta et al., 2012). The presence of this protein was later confirmed in guinea pig sperm, with a band of 120 kDa (Cordero-Martínez et al., 2018), which would partially agree with our results. Regarding ANO2, we observed a clear band at around 75 kDa, and another one at 150 kDa. As blocking peptide experiments demonstrated the specificity of the primary antibody, it could be that the lighter band corresponds to a degraded form of the protein, and the heavier band corresponds to a glycosylated form, given that this protein has multiple glycosylation sites. As no previous studies investigated the presence of ANO2 in mammalian sperm, no comparisons could be made.

Immunofluorescence revealed a distinct localization of ClCs and CaCCs. In effect, ClC-1 was localized in the post-acrosomal region and equatorial segment of sperm, ClC-2 was present in the tail, but the signal was more intense in the connecting and principal pieces than in the mid-piece. ClC-K was detected in the acrosome and the connecting and principal pieces. As none of these ClCs (i.e., ClC-1, ClC-2, and ClC-K) were immunolocalized in mammalian sperm, no comparisons between species could be made. It is, however, worth noting that ClC-3 was previously identified in the connecting piece and tail of sperm, which would agree with our results (Liu et al., 2017; Yeung et al., 2005). Regarding CaCCs, ANO1 was found to be distributed mainly along the sperm tail, with stronger fluorescence observed in the principal piece and discrete staining in the head region, but was absent from some sperm cells. This localization would partially agree with that of guinea pig sperm, as reported by Cordero-Martínez et al., who identified this channel in the mid-piece and apical region of the head (Cordero-Martínez et al., 2018). In our case, however, we did not observe that intense staining in the head and mid-piece; this could suggest species-specific differences in the localization of ANO1 in mammalian sperm. On the other hand, ANO2 was mainly found in the tail, with stronger staining in the mid-piece and some staining in the post-acrosomal sheath region of the head. As with the other ClCs investigated, no prior research has addressed the presence of ANO2 in mammalian sperm; thus, comparisons with other species could not be established. Despite this, it is worth emphasizing that our work demonstrates that members of the ClC and CaCC families are present in mammalian sperm, with distinct localizations across species and a potentially distinct role that warrants further research in species other than the pig.

After confirming the presence and localization of various ClCs and CaCCs in sperm, we aimed to investigate their role during sperm capacitation and acrosome reaction using three different pharmacological inhibitors: 9AC to block ClCs [32], BBR to inhibit CaCCs [50], and NPPB to block chloride channels in general. Based on these results, we then ran an additional series of experiments, in which BBR and NPPB were combined. Unfortunately, 9-AC and BBR do not specifically inhibit one type of channel (i.e., ClC-1/ClC-2, or ANO1/ANO2), but may block the entire family (e.g., ClCs), and can even impair the function of other channels, as it often occurs in research involving inhibitors in this and other cellular models. In the case of NPPB - and this is the reason it was chosen for testing - it inhibits CaCCs, as previously reported in the Wistar rat urinary bladder (Bijos et al., 2014), but also ClCs in retinal pigment epithelial (RPE) cells [35] and human sperm [36], and other chloride channels [51], including CFTR (H. Li et al., 2004).

Blocking CaCCs rather than ClC altered sperm motility, an effect that was observed after adding progesterone to induce the acrosome reaction. In effect, inhibiting CaCCs with BBR decreased total and progressive motility after 130 and 180 min of incubation, an effect that was also observed when sperm were treated with NPPB (200 µM), or a combination of BBR and NPPB (100 µM NPPB + 100 µM BBR). These results are consistent with a previous study conducted in guinea pigs, in which inhibiting ANO1 with T16Ainh-A01 and niflumic acid (NFA) reduced progressive and total motility after 120 min of incubation in Tyrode’s medium (Cordero-Martínez et al., 2018). In addition to these alterations in total and progressive motility, blocking CaCCs with BBR (100 µM) decreased VSL and VAP, an effect that was also observed when inhibiting chloride channels in general with NPPB (200 µM), but not when ClCs were blocked with 9AC. Furthermore, at the highest tested concentrations, BBR significantly reduced STR at 130 and 180 min of incubation. This decrease was also observed for the general inhibitor NPPB, which also decreased this variable at 120 min. Moreover, general inhibition of chloride channels with NPPB, but not specific blockade of CaCCs with BBR or ClCs with 9AC, significantly decreased ALH at 180 min, suggesting that regulation of ALH involves other chloride channels. Finally, BCF was also reduced when CaCCs were inhibited with BBR. All these findings suggest that CaCCs, rather than ClCs, regulate sperm motility and kinematics during capacitation. Hyperactivated motility is a feature of capacitation that enables sperm to propel through the viscous oviductal environment [53,54], detach from the oviductal reservoir, and penetrate the oocyte vestments [54]. Earlier research identified this hyperactivated motility as being characterized by high VCL and ALH along with low LIN and WOB [55]. As we observed that blocking CaCCs led to a reduction of VSL, we suggest that these channels, rather than ClCs, may be involved in the mechanisms underlying the acquisition of hyperactivated motility. Yet, the fact that some additional effects from chloride channel inhibition with NPPB were observed, like in the case of ALH, indicated that chloride channels other than ClCs and CaCCs could also play a role.

Mitochondria are understood to be essential for the regulation of sperm motility and, in this study and, consistent with this relationship, the inhibition of CaCCs with BBR or that of chloride channels as a whole with NPPB or NPPB+BBR, but not blocking ClCs with 9AC, significantly decreased mitochondrial membrane potential in viable sperm. These data align with previous research in human colon HT-29 cancer cells, where inhibiting CaCCs with CaCCinh-A01 also reduced mitochondrial membrane potential, decreased the BCL-2/BAX ratio, and subsequently triggered apoptosis through caspase 9 activation [56]. The decrease in mitochondrial membrane potential could be related to changes in superoxide levels, as an increase in these chemical species was observed in sperm treated with 100 µM BBR after 180 min of incubation. Because total ROS levels did not change, these findings would indicate that CaCCs inhibition with BBR causes localized mitochondrial oxidative stress, rather than generalized cellular oxidative stress, as superoxides are mainly produced in mitochondria [57,58]. Despite this, as neither the other concentrations of NPPB nor the combination of NPPB and BBR, which both reduced mitochondrial membrane potential, did not induce variations in superoxide levels, additional research is required to understand whether there is a cause-and-effect relationship between mitochondrial membrane potential and superoxide levels when CaCCs are inhibited, and why this occurs for BBR but not for NPPB. The fact that NPPB inhibits other chloride channels might provide a partial, but not an entire, explanation.

Inhibiting ClCs with 9AC increased membrane lipid disorder after 60, 120, and 180 min of incubation. This effect, however, was observed only in the total but not in the viable sperm population, which would indicate that membrane lipid disorder was rather higher in non-viable sperm than in their viable counterparts. Supporting these findings, the percentage of viable sperm with low membrane lipid disorder decreased following inhibition of ClCs, and that of non-viable sperm with high membrane lipid disorder increased (Suppl. Fig. 2a). On the other hand, while inhibiting CaCCs with BBR, or treating sperm with NPPB did not have any effect, combining NPPB and BBR (100 µM NPPB + 100 µM BBR, and 200 µM NPPB + 100 µM BBR) increased membrane lipid disorder, yet this effect was only observed after 60 min of incubation. This impact could be explained by the ability of NPPB to block other channels, including ClCs. Together, these results suggest that, in contrast to what is observed for motility and mitochondrial membrane potential, ClCs may be involved in regulating membrane lipid disorder during sperm capacitation and progesterone-induced acrosome exocytosis, whereas CaCCs contribute less. To the best of our knowledge, this is the first study interrogating the effects of inhibiting ClCs and CaCCs on membrane lipid disorder. For this reason, further research is needed to fully elucidate the relationship between chloride channels and the capacitation events that lead to cholesterol efflux and increase membrane fluidity.

Blocking CaCCs with BBR and BBR+NPPB significantly reduced tyrosine phosphorylation at 60 min and 180 min of incubation, respectively, an effect not observed when ClCs were blocked with 9AC. These results are in agreement with a previous study in guinea pigs, where incubating sperm in a chloride-deficient medium or in the presence of ANO1 inhibitors (T16Ainh-A01 and NFA) significantly reduced tyrosine phosphorylation; an effect that was already observed at 0 min and persisted after 60 and 90 min of incubation [59]. In contrast to this, neither acrosome integrity nor intracellular calcium levels were altered when ClCs and CaCCs were inhibited. In the case of NPPB, which also blocks ClCs, the lack of an effect on acrosome integrity would be in agreement with Roa-Espitia et al. (2025), who did not observe significant effects from inhibiting ClC-3 on the acrosome integrity of guinea pig sperm after incubation in Tyrode’s HEPES medium at 37 °C for 60 min (Roa-Espitia et al., 2025). In contrast, our findings would not align with those of Cordero-Martínez et al. (2018) and Roa-Espitia et al. (2025), who found that inhibiting ANO1 with T16Ainh results in a reduction of intracellular calcium levels and acrosomal integrity. While differences between species could explain these differences, further research is needed to confirm our findings.

## 5. Conclusions

In conclusion, we have demonstrated that ClCs and CaCCs are present in mammalian sperm, with distinct localizations that may be channel- and species-specific. We also observed that they are both involved in regulating sperm capacitation and acrosome exocytosis, with CaCCs appearing to play a more important role than voltage-gated chloride channels (ClCs), as inhibiting them disrupts key processes such as motility, mitochondrial function, and tyrosine phosphorylation. The role of ClCs appears more limited, yet they seem to regulate the increase in membrane lipid disorder that occurs during capacitation. Further research in species other than the pig is warranted to address whether their function is evolutionarily conserved across eutherian mammals.

## Declarations

### Ethics approval and consent to participate

Not applicable. Animals were handled exclusively by farm staff in accordance with the guidelines established by the Regional Government of Catalonia (Barcelona, Spain). The authors were not involved in animal handling, and the seminal doses used were commercial products intended for artificial insemination. Hence, specific approval from an ethical committee was not required for this study.

### Consent for publication

Not applicable. All data have been produced by the authors.

### Availability of data and material

The raw data for this study are available at CORA (https://doi.org/10.34810/DATA3385).

### Author’s contributions

MY and EP conceived the study. AA, JMH, CC, AA, KNB, EP, and MY conducted laboratory analysis. AA, EP, and MY discussed the results. AA drafted the Manuscript. MY and EP revised and edited the Manuscript. All authors read and approved the final version of the Manuscript.

## Acknowledgements

Not applicable.

## Use of AI

The authors did not use artificial intelligence tools in this study.

